# How osteons form: A quantitative hypothesis-testing analysis of cortical pore filling and wall asymmetry

**DOI:** 10.1101/2023.10.26.564273

**Authors:** Solene G. D. Hegarty-Cremer, Xenia G. Borggaard, Christina M. Andreasen, Bram C. J. van der Eerden, Matthew J. Simpson, Thomas L. Andersen, Pascal R. Buenzli

## Abstract

Osteon morphology provides valuable information about the interplay between different processes involved in bone remodelling. The correct quantitative interpretation of these morphological features is challenging due to the complexity of interactions between osteoblast behaviour, and the evolving geometry of cortical pores during pore closing. We present a combined experimental and mathematical modelling study to provide insights into bone formation mechanisms during cortical bone remodelling based on histological cross-sections of quiescent human osteons and hypothesis-testing analyses. We introduce wall thickness asymmetry as a measure of the local asymmetry of bone formation within an osteon and examine the frequency distribution of wall thickness asymmetry in cortical osteons from human iliac crest bone samples from women 16–78 years old. Our measurements show that most osteons possess some degree of asymmetry, and that the average degree of osteon asymmetry in cortical bone evolves with age. We then propose a comprehensive mathematical model of cortical pore filling that includes osteoblast secretory activity, osteoblast elimination, osteoblast embedment as osteocytes, and osteoblast crowding and redistribution along the bone surface. The mathematical model is first calibrated to symmetric osteon data, and then used to test three mechanisms of asymmetric wall formation against osteon data: (i) delays in the onset of infilling around the cement line; (ii) heterogeneous osteoblastogenesis around the bone perimeter; and (iii) heterogeneous osteoblast secretory rate around the bone perimeter. Our results suggest that wall thickness asymmetry due to off-centred Haversian pores within osteons, and that nonuniform lamellar thicknesses within osteons are important morphological features that can indicate the prevalence of specific asymmetry-generating mechanisms. This has significant implications for the study of disruptions of bone formation as it could indicate what biological bone formation processes may become disrupted with age or disease.

## 1. Introduction

Our skeleton is continuously renewed in order to preserve its strength and its fracture resistance by removing micro-damage and generating mineralisation heterogeneities that deflect crack propagation [1–3]. This renewal is performed by thousands of microscopic bone remodelling units (BRUs), that perform a tightly coupled and balanced resorption of old and damaged bone and subsequently infill the resorption cavities [4]. These BRUs include three phases: (i) an initial resorption phase, where osteoclasts perform the initial resorption guiding the path of the resorption cavities; (ii) a reversal-resorption phase, where osteoclasts enlarge the resorption cavities and where they are intermixed with osteoblastic reversal cells; and (iii) a formation phase, where osteoblasts infill the resorption cavity [5]. Each remodelling transaction results in a new bone structural unit (BSU). In cortical bone, BSUs are called secondary osteons or Haversian systems. The original resorption cavity is outlined by the cement line and the portion of the resorption cavity not infilled by bone formation remains as a Haversian canal [4, 6– 8].

Numerous histomorphometric studies of cortical osteons have provided insights into cortical bone remodelling during health, ageing, and disease [7, 9–18]. Most have focused on the mean dimensions of the osteons as a measure of the mean size of the resorption cavity (osteon area, osteon diameter), the mean magnitude of infilling (mean wall thickness), and the overall remodelling balance between these two activities (pore area, pore diameter). Much remains to be discovered about the unique characteristics and geometry of the individual osteons, shown to be highly variable [7, 19]. Osteons are the result of intracortical remodelling events either generating a new Haversian system (type I remodelling) or remodelling an existing Haversian system (type II remodelling). These different modes of remodelling affect the final morphology of osteons observed histologically [7, 19–25]. Drifting osteons have also been reported, resulting in osteons with distinctive morphologies, such as highly asymmetric wall thicknesses and off-centre pores [25–27]. Interpreting how different cellular processes involved in bone remodelling give rise to the observed osteon morphologies remains to be investigated.

Bone biopsies capture a single snapshot in time of the bone tissue remodelling state, but several cues are imprinted into the osteons during the bone formation processes, and can be analysed on biopsies using histomorphometry. Lamellae, osteocytes, mineral density, and mineralization labels provide valuable information related to bone formation variables. Measuring these quantities makes it possible in principle to backtrack some of the processes involved in osteon generation [28, 29]. However, there are complex interactions between cellular processes, geometric processes, and mechanical processes, which require careful consideration [4, 30–36]. Studies using dual mineralization labels have shown that matrix apposition rate (MAR) in osteons correlates strongly with resorption cavity radius [37–40]. This indicates a strong interplay between cellular processes—the production of new bone at a given time, represented by MAR—and geometry, represented by the cavity radius. In these studies, irregular (asymmetric) osteons were reported to have no well-defined cavity radius and are outliers in such relationships [37]. Several mathematical models have been developed to understand the influence of the geometry of bone surfaces on the rate of bone formation during pore filling [30, 32–36, 41, 42]. These cell population models elucidate the influence of the radius of curvature of the infilling pore on formation rate and match double labelling data in regular osteons very well [32, 34]. These mathematical models capture other qualitative features observed in osteon infilling, such as smoothing of initially irregular cement lines, and bone formation slowdown as infilling proceeds [34]. However, these models have not yet been quantitatively compared with detailed osteon data and none have investigated systematically how asymmetric osteons may be generated.

The frequency with which cortical osteons occur with a given degree of asymmetry, and how this frequency varies with age, are currently unknown. Understanding quantitatively how osteons may be generated with a given degree of wall thickness asymmetry is important to provide insights into how cortical pore filling may be regulated in health, and how this regulation evolves with age or disease. Our aims are therefore (i) to quantify the degree of osteon asymmetry in human cortical bone samples across ages, and (ii) to infer osteon formation mechanisms from observations of morphological features of osteons in bone samples. This requires quantitative considerations of the influence of cortical pore geometry on cellular processes during bone formation, which we base on a mechanistic mathematical model.

In this work, we first quantify osteon asymmetry based on differences in wall thickness around the osteon perimeter. We measure wall thickness asymmetry in human bone samples from women 16–78 years old and analyse the frequency distribution and the evolution with age of this measure of osteon asymmetry. We then use a mathematical model of cortical pore filling to simulate bone formation within osteon boundaries extracted from the human bone samples. Calibration of the mathematical model is performed based on osteons possessing a low degree of asymmetry. Various mechanisms for generating asymmetric osteons are incorporated into the mathematical model, including delayed initiation of bone formation in some region of the resorption cavity, differences in osteoblastogenesis around the resorption cavity, and differences in osteoblast secretory rate around the resorption cavity. We systematically explore the implication of these possible mechanisms for the generation of asymmetric wall thicknesses in irregular osteons by comparing numerical simulations of the mathematical model with morphological features observed in asymmetric osteons from the bone samples.

## 2. Materials and Methods

### 2.1 Bone specimens

We selected 3084 quiescent intracortical osteons within cortical bone biopsies previously subjected to a detailed histological analysis by Andreasen and colleagues [7, 20]. Regular and irregular osteons were included to assess their bone formation asymmetry. The bone specimens were collected from the iliac crest of 35 women (aged 16–78 years) during a forensic examination due to a sudden unexpected death. None of the women were receiving any medication affecting bone metabolism nor were any of the subjects diagnosed with metabolic bone diseases according to their medical records [7, 20].

In short, these undecalcified biopsies were fixed in 70% ethanol, methylmethacrylate-embedded, cut into 7.5 μm-thick sections and Masson’s trichrome stained or osteopontin immunostained for imaging and detailed cortical histomorphometry as described in [7, 20]. The study was approved by the Medical Ethical Committee Erasmus MC (2016-391) in compliance with the World Medical Association Declaration of Helsinki-ethical Principle for Medical Research Involving Human Subjects.

### 2.2 Histomorphometry

The histomorphometric analysis focused on bone formation asymmetry within the osteons across all samples. For each quiescent osteon, bone formation asymmetry was estimated by calculating the wall thickness (W.Th) asymmetry ratio, defined as the ratio of the two opposite W.Th having the largest difference (Figure 1A):

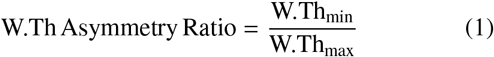

It is important to emphasise that W.Th asymmetry ratio is a measure of asymmetry of bone formation only. It does not quantify the degree of asymmetry of the overall appearance of the osteon and its resorption cavity. A W.Th asymmetry ratio value of one reflects an osteon in which the Haversian canal appears centred within the resorption cavity, however asymmetric this cavity may be, while a value of zero reflects an osteon in which no bone formation occurred on one side, such that the Haversian canal borders the cement line.

**Figure 1.**
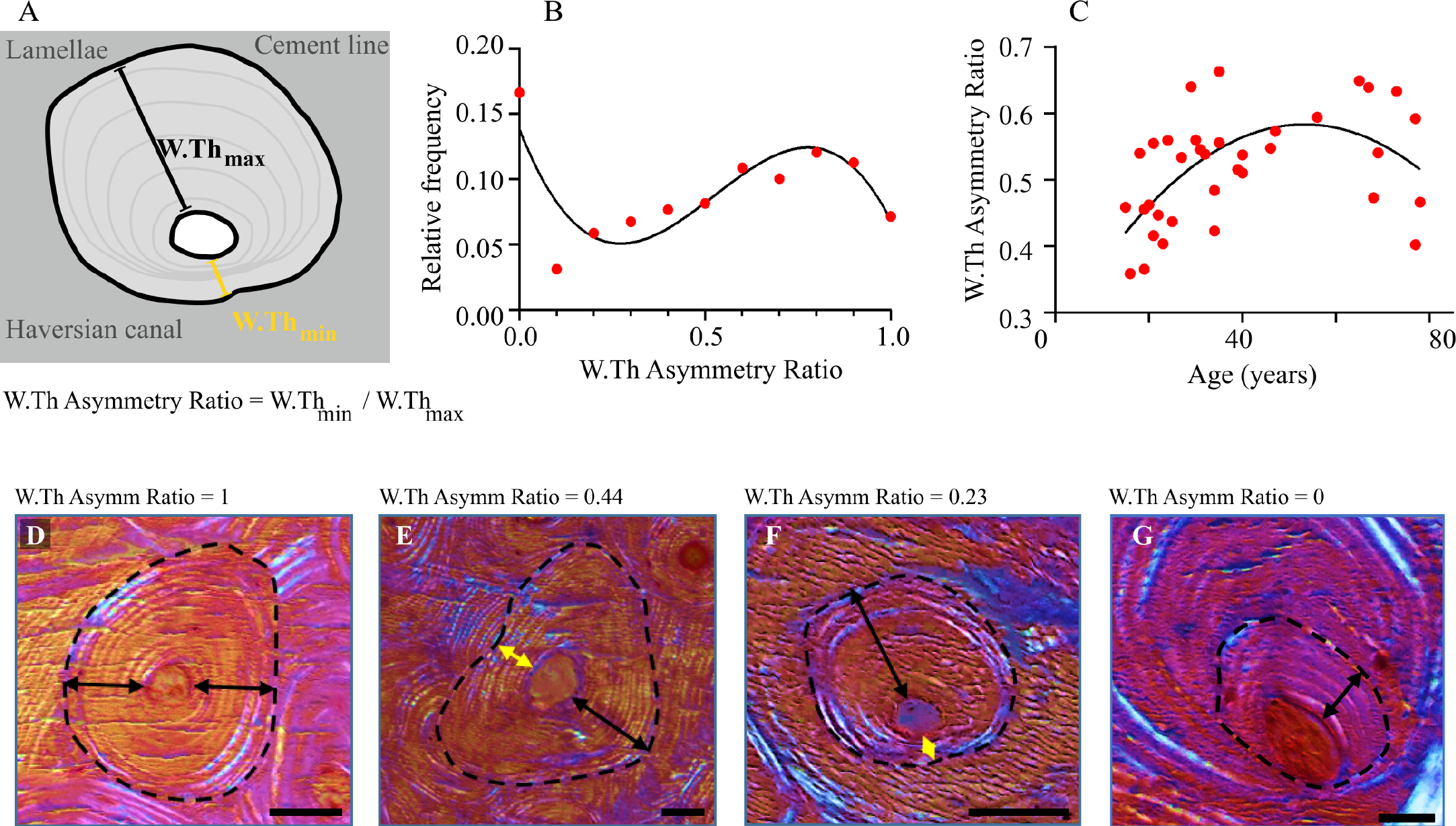
Osteon wall thickness asymmetry. (A) Wall thickness asymmetry is defined as the ratio of wall thicknesses on either side of the Haversian canal, in the direction in which asymmetry is highest; (B) Relative frequency of osteons with a given wall thickness asymmetry ratio in human bone samples 16–78 years of age; (C) Relationship between wall thickness asymmetry ratio and age in human bone samples, with a peak at 50–60 years (Spearman’s correlation analysis, p=0.01). Best fit second-order polynomial (black) W.Th.Asymmetry 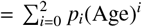 has coefficients *p*_0_ = 0.2666, 95% CI [0.12520, 0.408], *p*_1_ = 0.01189, 95% CI [0.00465, 0.0191], *p*_2_ = −0.0001116, 95% CI [− 0.000188, −0.000035] (D) Osteon with wall thickness asymmetry ratio of 1; (E) Osteon with wall thickness asymmetry ratio of 0.44; (F) Osteon with wall thickness asymmetry ratio of 0.23; (G) Osteon with wall thickness asymmetry ratio of 0. Scale bars in (D-G): 50 °m.

### 2.3 Mathematical model

The mathematical model describes the joint evolution of the bone tissue surface and of the density of osteoblasts on the bone surface. This model is based on previous mathematical models of tissue growth [34, 36] and is suitably extended to describe the infilling of symmetric and asymmetric osteons. The model includes the crowding or spreading of osteoblasts as the bone surface evolves, due to changes in local surface area, as well as the tendency of osteoblasts to even out their density by diffusive or directed cell motion along the bone surface [34, 36]. The model also incorporates specific mechanisms by which asymmetric bone formation may occur around the osteon perimeter, and an elaborate calibration procedure is proposed based on real osteon data. In the following, we present a non-technical summary of the assumptions of the mathematical model. Table 1 lists the mathematical symbols used in the model and how they relate to experimental observations. Further details about the mathematical model and its numerical simulation are presented in Appendix A.

**Table 1.** Mathematical symbols.

| Symbol | Description |
| --- | --- |
| $\rho$ | Osteoblast density [cells mm<sup>-1</sup>]. Number of osteoblasts per unit bone perimeter in a transverse cross section (N.Ob/B.Pm). The density of osteoblasts at the onset of formation is assumed to be $\rho_0 = 161$ cells mm<sup>-1</sup> [34], and the reference density of bone lining cells at the end of formation is assumed to be $\rho_{\text{end}} = 46$ cells mm<sup>-1</sup> [34, 43]. |
| $k_f$ | Secretory rate [mm<sup>2</sup>day<sup>-1</sup>]. Area of new bone secreted per osteoblast per unit time in a transverse cross section. Given by Eq. (11) with $a_k = 3.2741 \times 10^{-6}$ mm<sup>2</sup>day<sup>-1</sup> and $b_k = 8.5728 \times 10^{-5}$ mm<sup>2</sup>day<sup>-1</sup> [34]. |
| $k_f \rho = \text{MAR}$ | Local speed of the bone surface [mm day<sup>-1</sup>]. Corresponds to Mineral Apposition Rate (MAR) if there is no mineralisation defect. |
| $D$ | Diffusivity [mm<sup>2</sup> day<sup>-1</sup>]. Modulates the rate at which osteoblasts tend to even out their spatial distribution along the bone perimeter (not assessable by histomorphometry). |
| $A_0$ | Osteoblast elimination rate [mm day<sup>-1</sup>]. Modulates the rate at which active osteoblasts are eliminated, which captures the balance between osteoblast apoptosis, anoikis, and proliferation. This is assessable experimentally as the apoptosis index minus osteoblast proliferation and recruitment index. |
| $k_f \text{Ot}$ | Osteoblast embedment rate [day<sup>-1</sup>]: describes the rate at which osteoblasts are embedded into bone matrix to generate an osteocyte density Ot [cells mm<sup>-2</sup>] in a transverse cross-section, assumed to be $\text{Ot} \approx 625$ cells mm<sup>-2</sup> [34]. |

Osteon infilling is modelled by considering a two-dimensional transverse cross-section of the osteon, and the density of osteoblasts along the bone-cavity interface, denoted by *ρ* (number of osteoblasts per unit length of the bone perimeter), see Figure 2C. The initial shape of the bone-cavity interface assumed in the mathematical model at the onset of osteon infilling is the cement line of osteons identified from stained histological cross-sections (Figure 2A,B).

**Figure 2.**
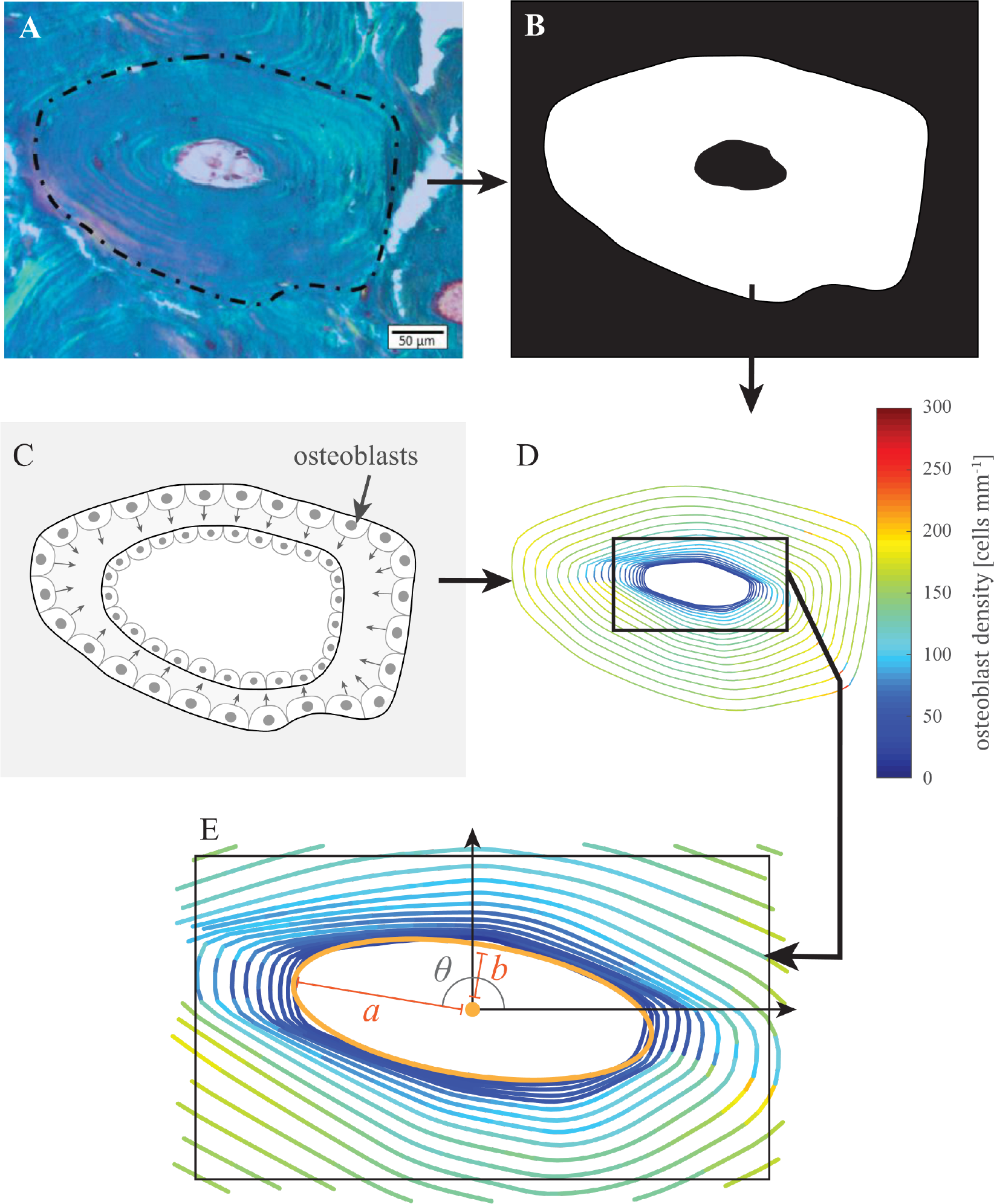
Image processing of experimental data and workflow for the simulation and calibration of the mathematical model. Cement lines and Haversian canals are identified in histological images of osteons (A). They provide a binarised representation of newly formed bone for use in the mathematical model (B). The mathematical model uses the identified cement line as initial bone surface for the population of osteoblasts (C) which produces new bone until pore area matches the area of the experimentally determined Haversian pore (C,D). The comparison between the location and shape of the simulated and actual Haversian pore is used to calibrate the mathematical model (E).

An initial population of osteoblasts is assumed to be present along the osteon boundary at the onset of bone formation [5]. As the cavity fills in, this population of osteoblasts evolves in time due to the balance of cell proliferation, cell death, cell detachment from the surface, and embedment into the bone matrix as osteocytes, leading to an overall depletion of actively secreting cells [34, 44]. For symmetric osteons, the initial population of osteoblasts is assumed to occupy the pore perimeter evenly with an initial osteoblast density of *ρ*_0_ ≈ 161 cells mm^−1^ [32, 34, 43]. As new bone tissue is deposited, the bone surface, and the osteoblasts that lie on it, progress into the resorption cavity in the cross section. Osteoblasts are assumed to produce new bone tissue at a rate *k*_f_ (area of bone formed per cell per unit time), and to be displaced perpendicularly to the underlying interface by this process (Figure 2C). The displacement of osteoblasts in the perpendicular direction crowds or spreads osteoblasts depending on the local geometry of the bone surface, such as its curvature, which generates spatial heterogeneities in osteoblast density [33, 34, 36].

The elimination of actively secreting osteoblasts is governed by a first-order reaction rate *A*, which determines how quickly the osteoblast population depletes due to the balance between cell proliferation, cell death (e.g., by apoptosis) and detachment from the bone surface (anoikis). The strength of osteoblast elimination rate is governed by the parameter *A*_0_ (Appendix A, Eqs (9)–(10)). The embedment of osteoblasts into the bone matrix and their differention into osteocytes similarly depletes the population of active osteoblasts with a rate given by Ot × MAR [28, 29, 34].

The diffusive motion of osteoblasts along the interface describes the tendency of osteoblasts to even out their density due to local cell-cell mechanobiological interactions [45–48]. This diffusive redistribution of osteoblasts counters heterogeneous crowding effects induced by the geometric changes in available bone surface area during infilling. In the model, the rate of diffusion is governed by the diffusivity parameter *D*. The larger this parameter *D*, the more quickly osteoblasts return to a uniform density.

By including geometric cell crowding effects, cell diffusion, cell secretory rate, cell proliferation, cell elimination rate, and differentiation into osteocytes, the mathematical model describes changes in osteoblast density around the interface over time, and changes in the bone deposition rate around the bone pore perimeter (Figure 2D; Appendix A, Eqs (8)–(16)).

Numerical simulations of osteon infilling are performed starting from the outline of resorption cavities (cement line) visible in the experimental images (Figure 2A,B). Infilling is assumed to continue until the pore area in the simulation reaches the area of the Haversian canal from the experimental image, or until the density of osteoblasts reaches a minimum value, the density of bone lining cells, *ρ*_end_ ≈ 46 osteoblasts mm^−1^ [34, 43].

To simulate the infilling of osteons possessing strong degrees of asymmetry of wall thickness, further mechanisms are incorporated into the model. Based on inspections of histological cross-sections, three hypotheses about further processes that may generate strong osteon wall asymmetries are considered:

H1. A delay in the initiation of bone formation on a portion of the cement line;

H2. An uneven generation of osteoblasts which causes faster infilling in some region of the bone pore perimeter;

H3. An uneven cell secretory rate, which also cause faster infilling in some region of the bone pore perimeter.

Hypothesis H1 is modelled by setting the secretory rate *k*_f_ to 0 on a certain portion of the cement line for a set amount of time at the beginning of the simulation (Appendix A, Eq. (12)–(14)). This will result in that section of the bone pore perimeter commencing infilling later. Hypothesis H2 is modelled by adjusting the initial density of osteoblasts around the bone perimeter. Sinusoidal perturbations of density around the perimeter are chosen, such that the total count of osteoblasts at the start of infilling remains the same (Appendix A, Eq. (15)). Hypothesis H3 is modelled similarly to Hypothesis H2. A sinusoidal perturbation is added to the osteoblasts’ tissue secretion rate such that there is a region of the cortical pore with low secretory rate and a region of the cortical pore with high secretory rate, and the mean secretory rate around the interface remains the same (Appendix A, Eq. (16)).

The consequences of these hypotheses for generating asymmetric osteon walls are explored systematically and compared with osteons possessing strong degrees of wall thickness asymmetries.

### 2.4 Error metrics and calibration procedure

In our previous work [34], calibration procedures were based on optimising the circularity of Haversian canals at the end of simulations of infilling in artificially generated osteon shapes. Here we develop a more detailed calibration procedure based on matching simulated Haversian canal position and Haversian canal shape with experimental Haversian canals extracted from histological cross sections.

To calibrate the mathematical model, the diffusivity parameter *D* and elimination rate parameter *A*_0_ are systematically varied to find the combination which produces simulation results closest to the experimental data. These two parameters are chosen as they are the parameters with the most uncertainty experimentally, and they significantly affect the geometry of the simulated osteon. The discrepancy between simulation results and experimental data is quantified using a combination of five error metrics. Sampling these error metrics by running the model with different parameter values provides error surfaces closely related to parameter likelihood functions [49]:

1. *Final osteoblast density error* the relative error between the average density of osteoblasts around the bone perimeter at the end of the simulation and the expected density of osteoblasts at the end of infilling (bone lining cell density), *ρ*_end_ ≈ 46 osteoblasts mm^−1^ [34, 43],

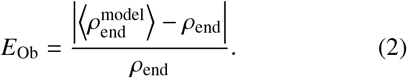
2. *Final pore area error:* the relative error between the area of the Haversian canal generated by the mathematical model and that of the experimental image (inner black area in Figure 2B),

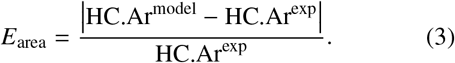
3. *Final pore location error:* the relative error between the location of the centroid of the Haversian pore, and the location of the centroid of the final pore from the simulation, normalised by the average diameter of the initial pore:

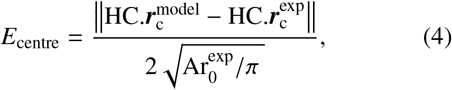

where 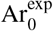 is the initial pore area.
4. *Final pore shape error:* the relative error in orientation and in aspect ratio of ellipses fitted to the final pore shape in the simulation and the experimental image (Figure 2E). The fitting procedure provides the ellipses’ semi-axes *a* and *b*, and the angle *θ* between the ellipses’ major axis and the *x* coordinate axis (in radians) (Appendix B). The relative errors in orientation *θ* and in aspect ratio *a/b* of the ellipses are given by:

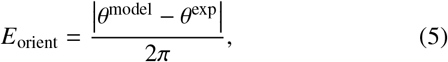

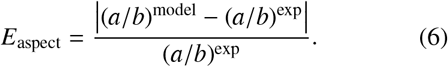

The error metrics on the area, location, orientation, and aspect ratio of the final pore correspond to a succession of approximations that involve the 0^th^, 1^st^, and 2^nd^ moments of area of the pore region (Appendix B). Most Haversian canals have a well-defined elliptic shape, so we did not consider higher order moments. The total error for simulation of infilling in a single osteon is the weighted sum of the above five error metrics:

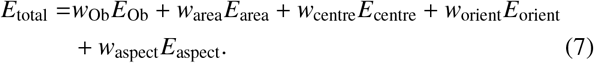

The total error for simulations of infilling of several osteons using identical model parameters is the sum of the total errors for each osteon.

### 2.5 Numerical simulations

The general workflow for numerical simulations of infilling is summarised in Figure 2:

1. The cement line and Haversian pore are first extracted manually from the experimental image of an osteon and digitalised into a mask (Figure 2A,B).
2. The digitalised cement line provides the initial resorption cavity within which bone infilling is simulated (Figure 2C,D)
3. The mathematical model is solved numerically by assuming an initial population of osteoblasts along the walls of the initial resorption cavity, fixed values for the diffusivity and elimination rate parameters *D* and *A*_0_, and a combination of asymmetry-generating hypotheses H1, H2, H3 (Figure 2D).
4. The infilling simulation is stopped when the remaining pore area matches the Haversian canal area from the experimental image, or when osteblast density falls below *ρ*_end_ (Figure 2D).
5. Quantitative comparisons between the simulated osteon and the experimental image are performed based on the error metrics in Eqs (2)–(7) (Figure 2E).

More detail on how the mathematical model is solved numerically is found in Appendix A.

## 3. Results

### 3.1 Wall thickness asymmetry of quiescent osteons within human cortical bone

The degree of infilling asymmetry assessed by the W.Th asymmetry ratio (Figure 1A, Eq. (1)) was highly variable among the 3084 osteons measured in the iliac crest of 35 women (Figure 1B). Osteons ranged from having completely symmetric pore infilling (W.Th asymmetry ratio = 1) to having completely asymmetric pore infilling with no bone formation on one side (W.Th asymmetry ratio = 0). The W.Th asymmetry ratio has a significant bell-shaped correlation with age with a peak at 50–60 years (p=0.01, Spearman’s correlation *r*_*s*_ = 0.44), meaning that W.Th asymmetry ratio increases until 50– 60 years of age after which it starts to decline (Figure 1C). Best fit second-order polynomial W.Th.Asymmetry 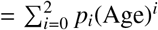 has coefficients *p*_0_ = 0.2666, 95% CI [0.12520, 0.408], *p*_1_ = 0.01189, 95% CI [0.00465, 0.0191], *p*_2_ = −0.0001116, 95% CI [−0.000188, −0.000035]. Examples of osteons with a range of W.Th asymmetry ratios are illustrated in Figures 1D–G.

### 3.2. Numerical simulations of symmetric osteon infilling

Figure 3 shows numerical simulations of pore filling for a single osteon with low bone formation asymmetry (Osteon G in Figure 5, W.Th asymmetry ratio = 0.8). The model is run without incorporating any asymmetry-generating processes H1–H3. Each panel in Figure 3 corresponds to a different combination of the diffusivity *D* and elimination rate *A*_0_. In each panel, the evolving bone interface is shown at twenty regularly spaced values of time by curves coloured by the density of osteoblasts, from time *t* = 0 until the final time of the simulation. For large values of *A*_0_, the total area of new bone formed is less than that in the bone sample. In this case, the simulation stops when the average osteoblast density reaches *ρ*_end_ ≈ 46 cells mm^−1^, leading to a large area error metric. For small values of *A*_0_, the total area of new bone formed matches precisely that from the bone sample since the simulation stops when this target area is reached. However, the final density of osteoblasts in the simulations,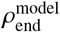, may be far from the expected final osteoblast density *ρ*_end_, resulting in a large density error metric.

**Figure 3.**
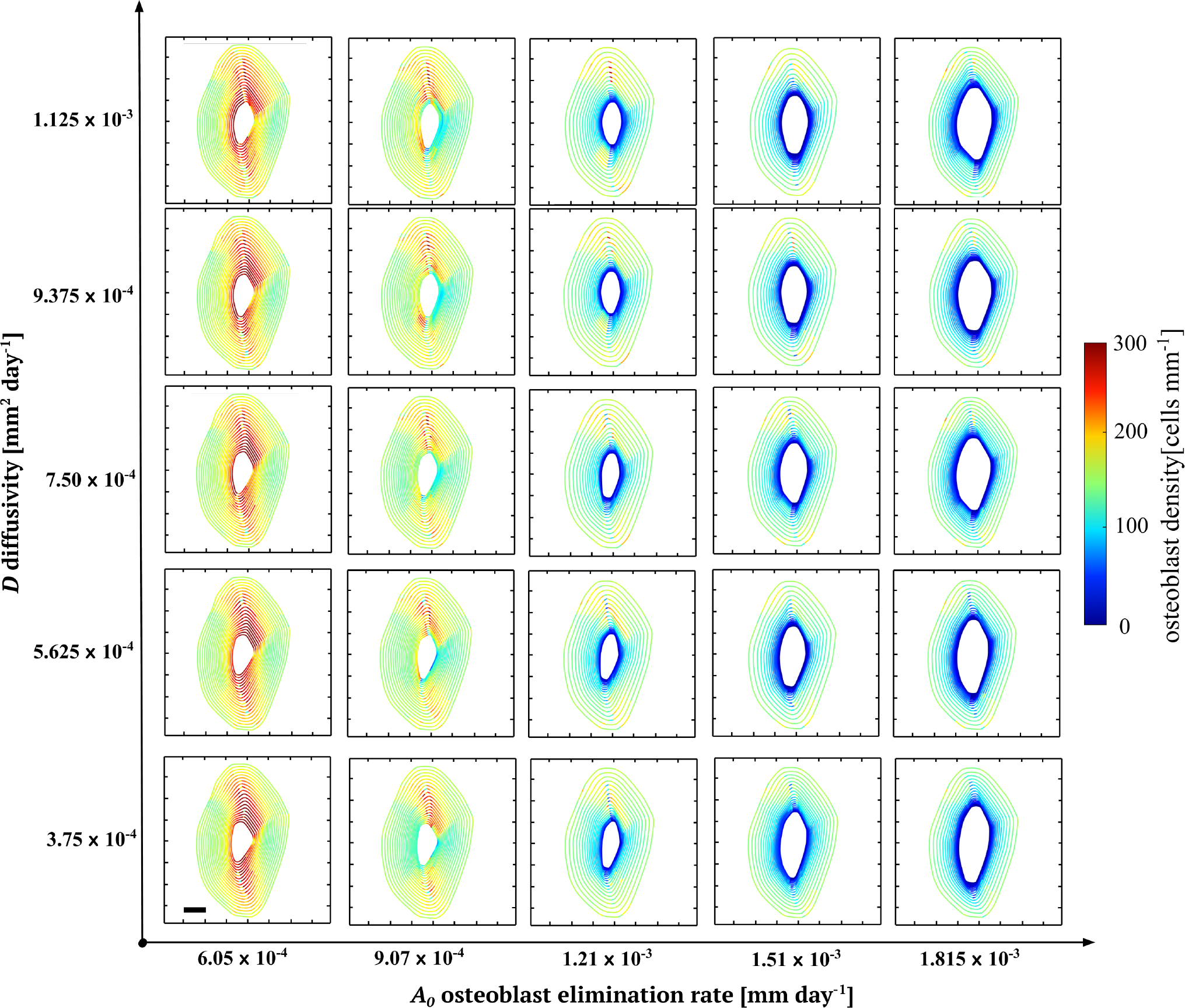
Influence of the diffusivity parameter *D* and osteoblast elimination rate parameter *A*_0_ of the mathematical model on osteon infilling. In each panel, the bone surface is shown at twenty regularly spaced values of time from *t* = 0, where it coincides with the experimental osteon boundary, until either the target final pore area is reached, i.e., the Haversian canal pore area measured on the experimental image, or until the density of osteoblasts is less than the reference bone lining cell density *ρ*_end_. A large elimination rate parameter *A*_0_ leads to a greater final pore area compared to the experimental image since the population of osteoblasts decreases to the threshold *ρ*_end_ before reaching the target pore area.

### 3.3. Calibration

Figure 4 cumulates specific and total error metrics obtained from numerical simulations of pore filling across a series of osteons with low formation asymmetry (W.Th Asymmetry Ratios between 0.525 and 0.9, shown in Figure 5). Since all the error metrics are relative errors, the total error metric in Figure 4A is calculated using equal weights in Eq. (7) (*w*_Ob_ = *w*_area_ = *w*_centre_ = *w*_orient_ = *w*_aspect_ = 1). The error metrics are sampled using combinations of *D* and *A*_0_ shown by black circles in panel A, and are interpolated between these combinations by MATLAB’s `contourf` function [50]. We can interpret these error metrics, shown as heat maps in Figure 4, as indication of the probability of particular parameter combinations given our fixed data [49].

**Figure 4.**
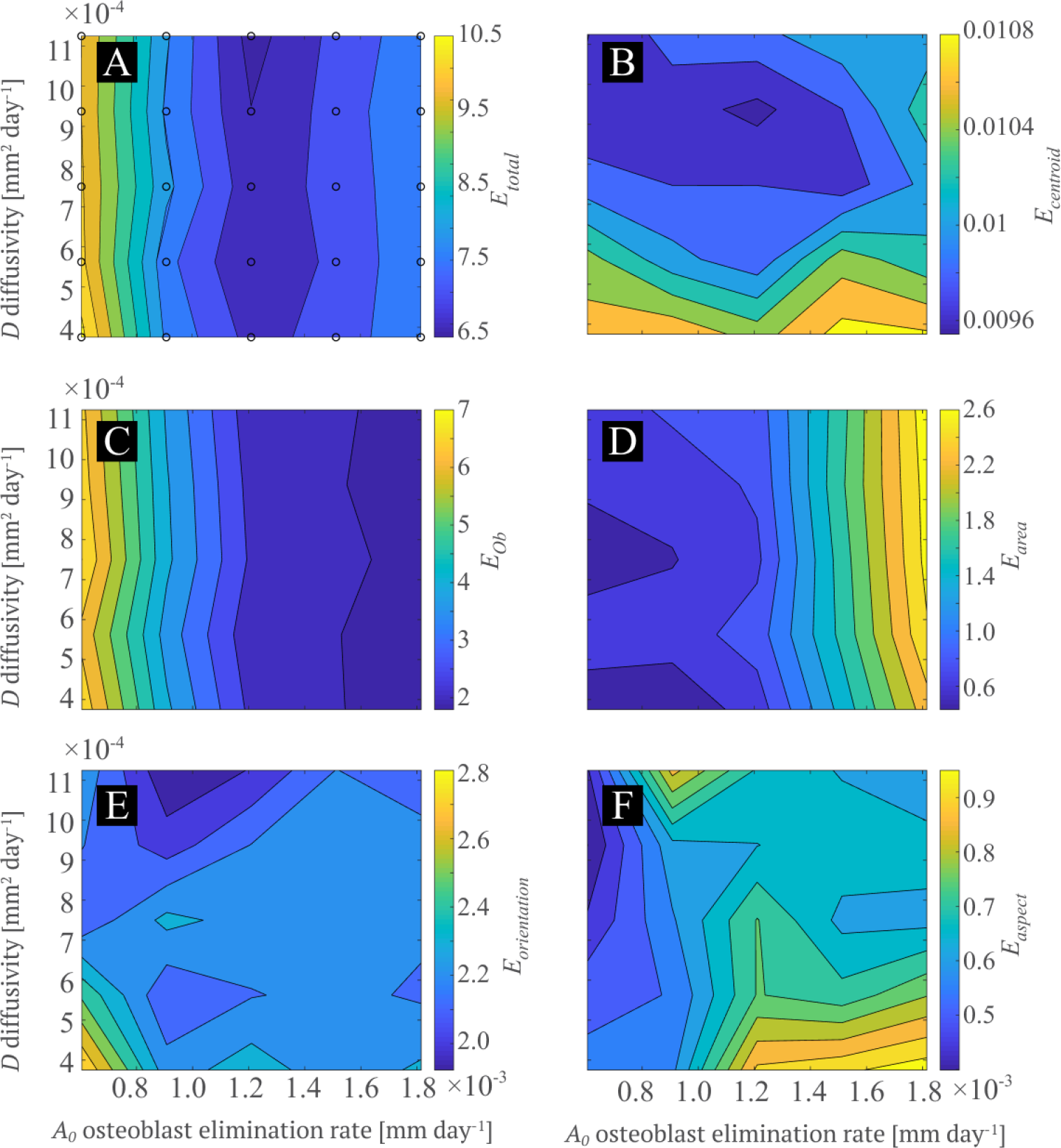
Parameter sweep and calibration of the mathematical model. (A) Total error between the biological data and the simulation results for each pair of *D* and *A*_0_ parameters calculated by Eq. (7) with *w*_Ob_ = *w*_area_ = *w*_centre_ = *w*_orient_ = *w*_aspect_ = 1. Each black circle represents a pair of parameters used in the numerical simulations to sample the total error. Error values at other points are interpolated by MATLAB’s function. The lowest total error occurs at *A*_0_ = 1.21 ×10^−3^ [mm day^−1^] and *D* = 1.125 ×10^−3^ [mm^2^day^−1^]. (B)–(F) Relative errors calculated from Eqs (2)–(6) at the same sampled parameter pairs (*D,A*_0_) shown in (A) and interpolated: centroid error (B), density error (C), area error (D), ellipse orientation error (E), ellipse aspect ratio error (F).

**Figure 5.**
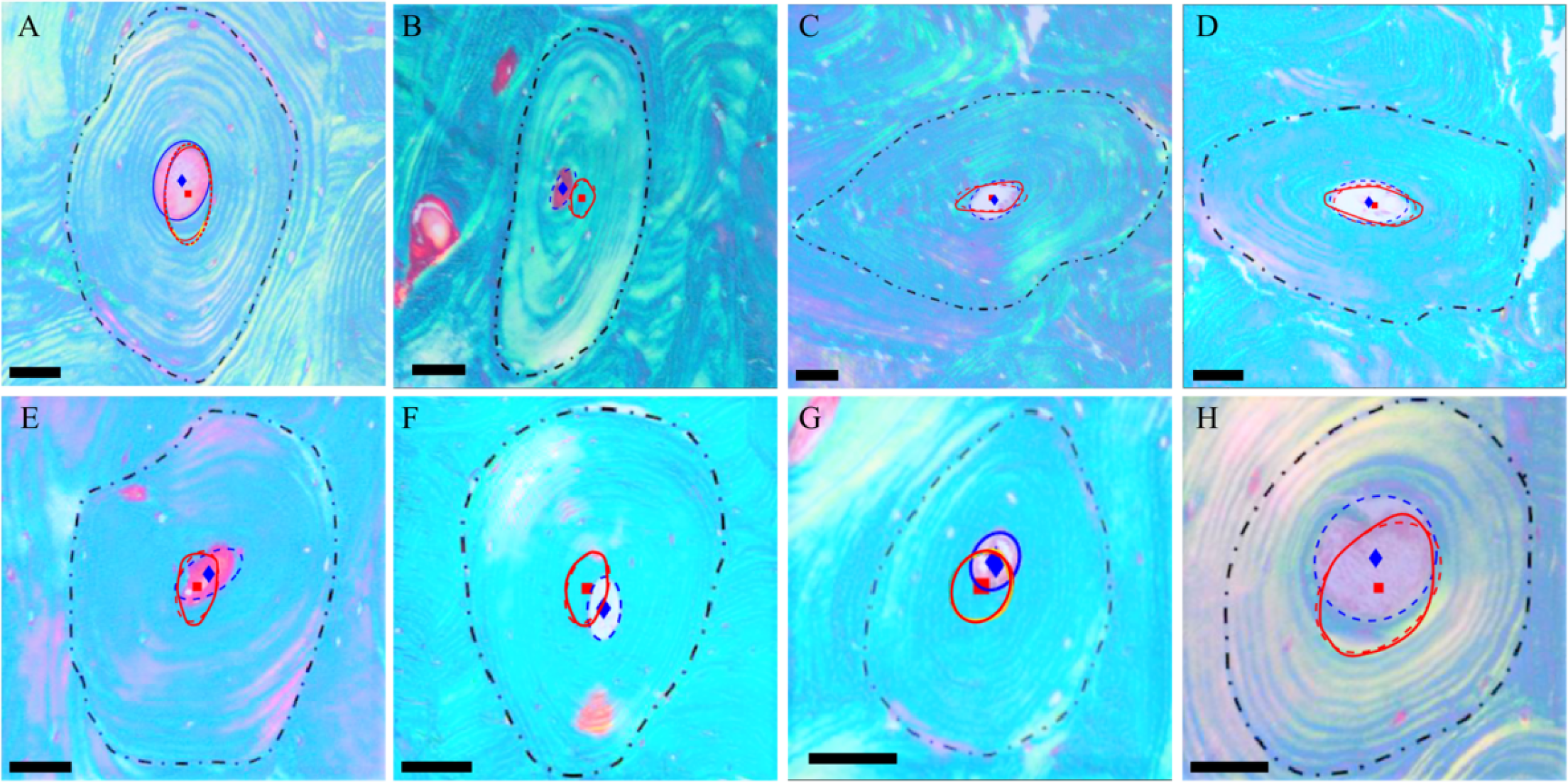
Osteons with low formation asymmetry (W.Th Asymmetry Ratios between 0.525 and 0.9) used in the parameter sweep to determine the optimal pair of parameters (*D, A*_0_). Blue diamonds highlight the centroid of the final pore in the biological data, and the blue ellipse shows the ellipse fit to the Haversian canal. Cement lines (dash-dotted black line) are used as initial bone surface for numerical simulations of infilling. Haversian canals calculated by the model using the optimal pair of parameters (*D* = 1.125 ×10^−3^ mm^2^*/*day, *A*_0_ = 1.21 ×10^−3^ mm day^−1^) are overlaid (solid red curves), as well as their elliptic fit (dashed red curves) and centroid (red square).

The density error metric (panel C) and area error (panel D) exhibit high sensitivity to the elimination rate *A*_0_, and weak sensitivity to osteoblast diffusivity *D*. The pore location error in panel B and, to some extent, the aspect ratio error in panel F exhibit more pronounced sensitivity to osteoblast diffusion than elimination rate.

With equal weighting of the individual relative errors, the total error metric is lowest at the middle value of *A*_0_ in these plots, *A*_0_ ≈ 1.211 ×10^−3^ mm day^−1^, and at *D*≈ 1.125 ×10^−3^ mm^2^ day^−1^. Other choices of weightings in the total error are possible and would change the parameter combination at which total error is lowest. However, Figure 5 shows that the final pores obtained by the model with these values of *A*_0_, *D* match well with the Haversian canals seen in the experimental images, despite strong variability in resorption cavity shapes. These values of *A*_0_ and *D* are therefore used as basis for testing hypotheses H1– H3 for asymmetric pore infilling in the next section.

### 3.4. Hypothesis testing of asymmetric osteon generation

In Figure 6, the three hypothesis H1–H3 formulated for asymmetric pore infilling are tested by simulating the infilling of an osteon (named Osteon A1) with intermediate bone formation asymmetry (W.Th Asymmetry Ratio = 0.55). This osteon possesses a slightly off-centred Haversian pore in the experimental image. All three hypotheses successfully displace the location of the final simulated pore from the centre of the resorption cavity toward its actual location in the experimental image. However, each hypothesis leads to distinctive intermediate evolutions of bone interface and osteoblast density that may be correlated with morphological features of the osteon.

**Figure 6.**
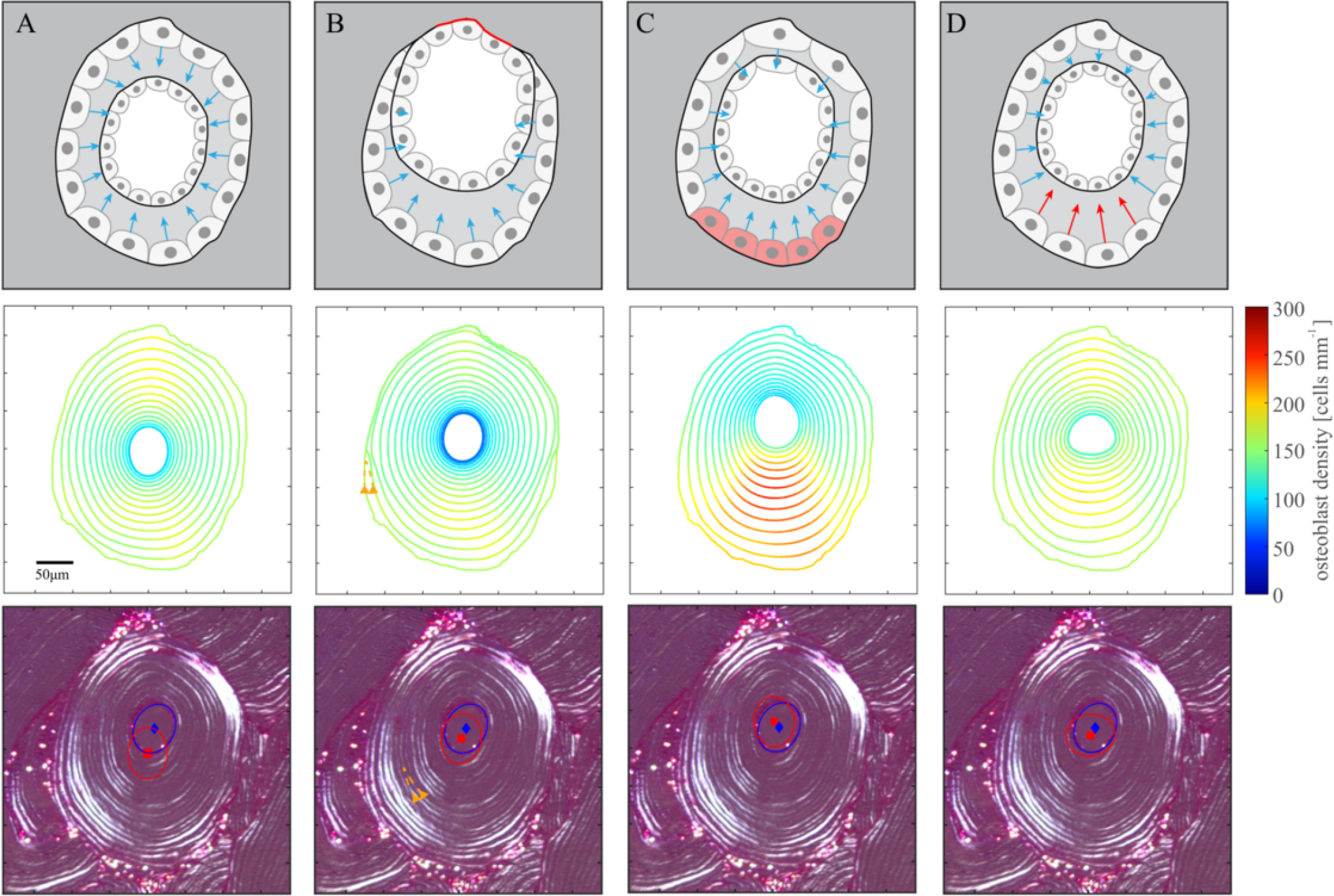
Results of simulations of asymmetric osteon A1 with an off-centred pore (W.Th Asymmetry Ratio = 0.55). The first row depict the asymmetry generating mechanism used in A–D schematically. The second row shows simulation results obtained, and the third row shows the experimental image overlaid with the ellipse fitted to the final pore of the simulation data (blue) and the ellipse fitted to the Haversian canal (red). Column A shows the results with no asymmetry generating mechanism. B shows the results under H1 (time delay), with an example of eclipsing lamellae highlighted in orange in both the biological data and the simulation, obtained with *T* = 10 days, *θ*_min_ = 0, *θ*_max_ = *π* in Eq. (14). C shows the results under H2 (nonuniform initial density of osteoblasts) with *c*_Ob_ = 0.5, *δ*_Ob_ = *π/*2 in Eq. (15). D shows the results under H3 (nonuniform secretory rate) with *c*_secr_ = 0.5, *δ*_secr_ = *π/*2 in Eq. (16).

Simulations performed by assuming delayed initiation of infilling in some region of the cement line (H1) show eclipsing lamellae behaviour, that is, a sequence of interfaces at different times that intersect in some part, and that enclose a crescent-shaped newly formed bone region (corresponding to a lamella) that does not circle around the osteon. This behaviour can be seen in the experimental image of Osteon A1 in Figure 6B (orange lines and arrow heads). Cell density around the final pore under hypothesis H1 is relatively homogeneous and close to the final density expected, *ρ*_end_.

Figures 6C,D show that when the initial speed of bone formation is heterogeneous around the cement line (hypotheses H2 and H3), simulations of infilling produce variations in thickness of individual lamellae. In Figure 6C (H2), a high density of osteoblasts can be seen for most of the simulation on the lower half of the osteon. Cell density results in Figure 6D (H3) are more homogeneous, however the orientation and aspect ratio of the final pore are less aligned with the experimental data.

Simulations of pore filling in a more asymmetric osteon (Osteon A2, W.Th Asymmetry Ratio = 0.26) are shown in Figure 7. For the model to reproduce the location and shape of the osteon more accurately, a combination of infilling asymmetry mechanisms is required. Figures 7D–F show that with a single asymmetry generating mechanism, the final pore may have a different shape, may not be offset enough compared to the experimental image (Figure 7D,E), or the density of osteblasts may be unreasonably high (Figure 7D,E,F). However, combining the mechanisms H1–H3 produces results closer to the experimental image (Figure 7B).

**Figure 7.**
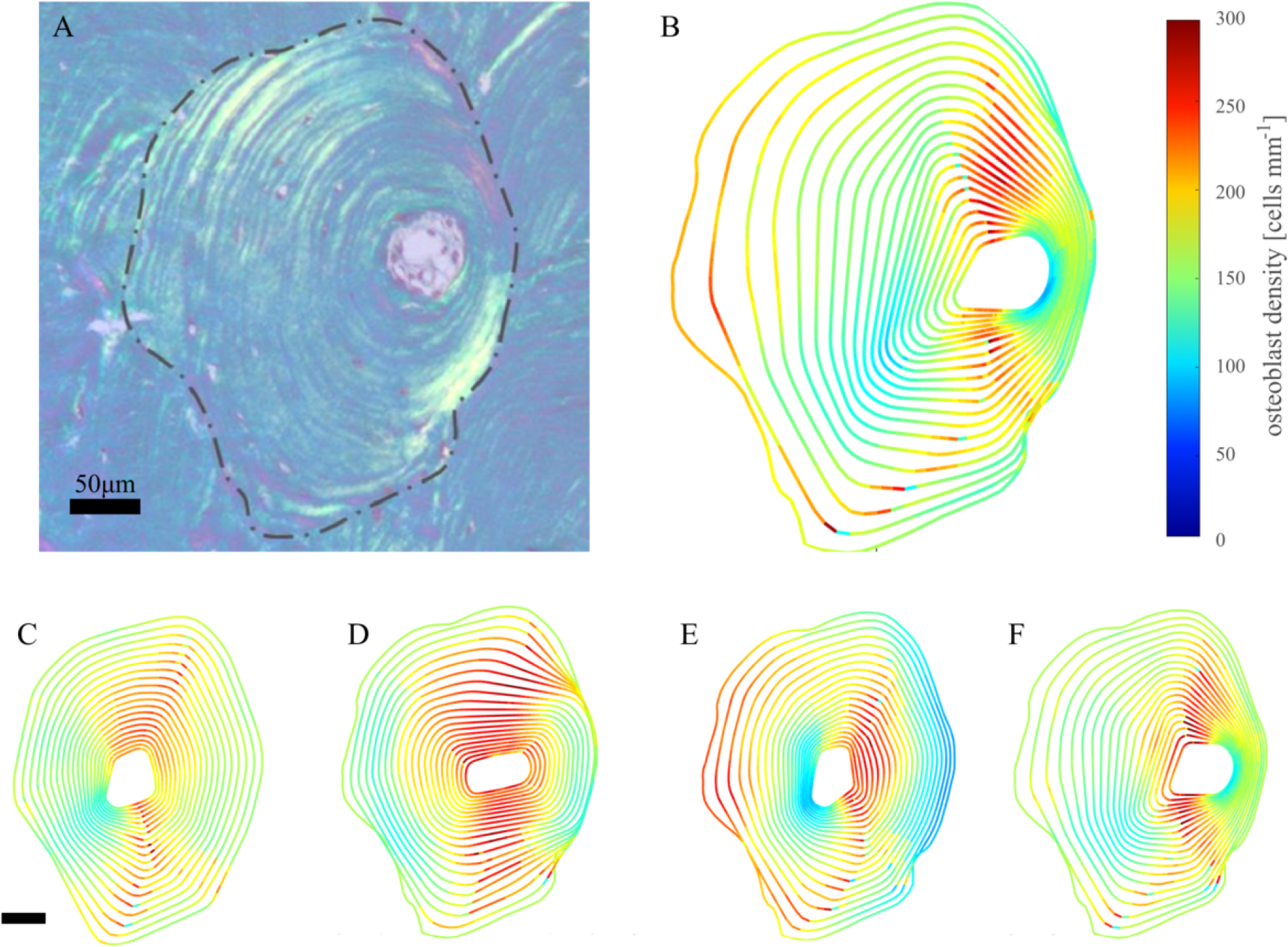
Results of simulation combining asymmetry generating mechanisms. A – Osteon A2, with W.Th Asymmetry Ratio = 0.26. B – Simulation result obtained by combining H1, H2, and H3, with *T* = 12, *θ*_min_ = −*π/*4, *θ*_max_ = *π/*4, *c*_Ob_ = 0.3, *c*_secr_ = 0.7, *δ*_Ob_ = 0, *δ*_secr_ = 0 in Eqs (12)–(16). C – Simulation result with no asymmetry generating mechanism. D – Simulation result under H1 with *T* = 25, *θ*_min_ = −*π/*4, *θ*_max_ = *π/*4 in Eq. (14). E – Simulation result under H2 with *c*_Ob_ = 0.5, *δ*_Ob_ = 0 in Eq. (15). F – Simulation result under H3 with *c*_secr_ = 0.7, *δ*_secr_ = 0 in Eq. (16).

W.Th Asymmetry Ratio is particularly sensitive to Haversian canal location within the resorption cavity. In Figure 8, we therefore summarise how model assumptions (*A*_0_, *D* and hypotheses H1–H3) affect final pore location error as a function of W.Th Asymmetry Ratio for osteons A–H of Figure 5 and osteons A1, A2 of Figures 6, 7.

**Figure 8.**
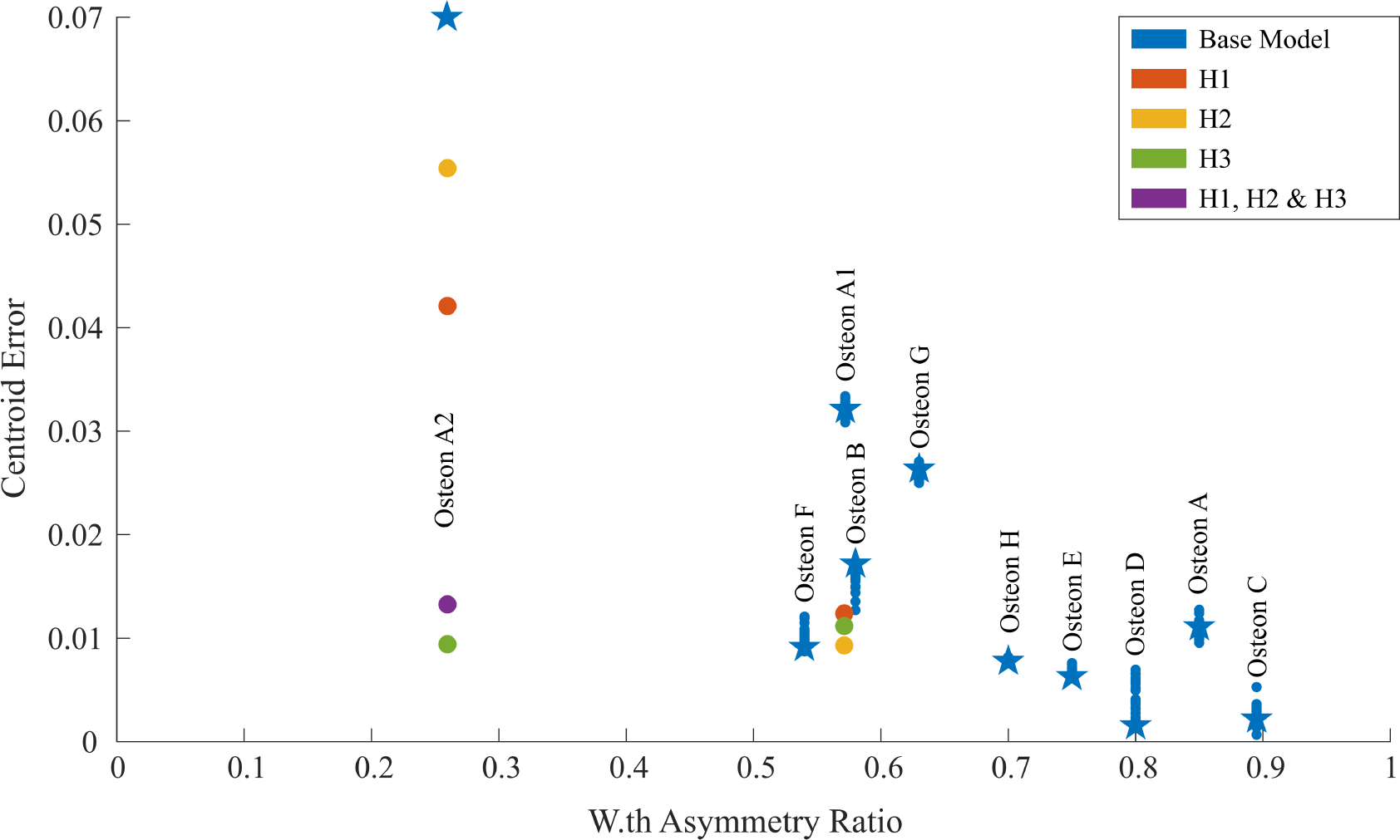
Centroid Error Metric (Eq. (4)) as a function of W.Th Asymmetry Ratio. The blue circular markers indicate centroid errors obtained by the parameter sweep without asymmetry generating mechanisms. The blue star markers show centroid errors for the optimal parameter pair as found by the parameter sweep. For osteons A1,A2, red markers indicate asymmetry generating mechanism H1 was added to the model, yellow markers indicate H2 was used, green markers indicate H3 was used, and purple markers indicate a combination of H1, H2 and H3 was used.

Without accounting for asymmetric infilling hypotheses H1– H3, pore location error increases for decreasing W.Th Asymmetry Ratios, as expected (Figure 8, blue markers). Pore location error is low for highly symmetric osteons, even with suboptimal diffusivity and elimination rate parameters (Osteons C,A,D,E,H). For low enough W.Th Asymmetry Ratios, no combination of model parameters *A*_0_, *D* can significantly reduce the centroid error metric, unless asymmetric infilling hypotheses are introduced (Osteons A1 and A2). We see that for Osteon A2, the largest pore location error is obtained without any asymmetry generating mechanism. The error decreases by considering single asymmetry generating mechanisms, and is lowest when H3 or all three mechanisms are combined.

## 4. Discussion

Quiescent osteons in human cortical bone are often depicted as a cylindrical structure with symmetric cutting cone and symmetric closing cone (W.Th asymmetry ratio = 1) [44, 51]. In reality osteons have a wide range of shapes, sizes and wall thickness asymmetry (Figure 1B). Osteons have been classified into several categories depending on their histomorphometric features, including type I ostons that create a new Haversian canal, type II osteons that remodel a previously existing vascular channel, and drifting osteons, in which resorption and formation occurs unilaterally for some time [5, 20, 26, 26, 52]. These different types of osteons arise from different modes by which bone formation occurs in resorption cavities, which can have a direct influence on W.Th asymmetry around the osteon. The irregularity of osteonal structures has often been reported based on serial cross-sections [21, 26, 52–54] and, more recently, micro-CT images [19, 55, 56]. However, few studies quantitatively assessed asymmetries of osteonal structures [19, 26].

Our measurements of W.Th asymmetry ratios indicate that perfectly symmetric osteons are rare. Most osteons analysed in our study possess some degree of asymmetry in wall thickness across the Haversian pore, and some are completely asymmetric, meaning that the formation process did not occur all around the resorption cavity. Osteons with a W.Th asymmetry ratio equal to zero or one are outliers in terms of frequency of occurrence in our study (Figure 1B), which suggests that they could be created under particular conditions. The prevalence of asymmetric osteons has a clear increasing trend with wall asymmetry ratio (Figure 1B), meaning that more symmetric osteon walls are more frequent. This non-normally distributed, increasing trend of prevalence with wall asymmetry ratio suggests the existence of systematic effects that influence the formation of new bone within osteons at a local level, and that could result in wall thickness asymmetries, such as mechanical gradients, proximity to neighbouring Haversian systems, or branching points of the Haversian canal structure. Our samples did not contain sufficient spatial information to determine possible correlations of wall asymmetry with skeletal sites and mechanics. Further studies would be required to investigate such links.

Figure 1C suggests that on average, W.Th asymmetry ratios also increase with age until about 50 years, and decreases thereafter. The increasing prevalence of more symmetric osteons from 20 to 50 years correlates with the gradual replacement of primary osteons created during growth by secondary osteons created by remodelling. The creation of more asymmetric osteon walls after 50 years of age could be due to age-related bone loss, and associated disruption of bone formation [57]. Bone formation tends to smooth irregular cavities, so a lack of bone formation could generate more asymmetric osteons [34].

Several processes influence how an osteon forms, including osteoblast recruitment, osteoblast elimination by anoikis or apoptosis, osteoblast activity (secretory rate) but also osteoblast lateral redistribution (diffusion) and the significant shrinkage of bone surface availability during pore infilling [30, 32, 34, 44]. Bone tissues record several dynamic processes during bone formation, but it is very challenging to link morphological features with the dynamic processes that created these features during bone formation experimentally [28, 29, 58]. Mathematical modelling can help provide such links and be used as a tool for hypothesis testing. We therefore formulated a mathematical model that includes the complex interplay between these processes, and calibrated the predictions of the model against experimental images of regular osteons. We then tested whether some morphologic features observed in histological cross sections of asymmetric osteons, such as wall asymmetries and eclipsing lamellae [59] could be related to changes in biological processes involved in cortical pore filling, namely: (i) delays in the onset of bone formation; (ii) changes in the distribution of osteoblasts around the resorption cavity; and (iii) spatial differences in osteoblast activity.

Uneven distributions of osteoblasts at the onset of formation leading to faster infilling on one side of the bone perimeter (Hypothesis H2) may arise due to uneven osteoblastogenesis. Osteoblastogenesis involves several cellular differentiation stages of pre-osteoblasts regulated by many growth factors and signalling molecules, such as Wnt, sclerostin, TGF*β*, PTH, Eph/Ephrin, and the RANK-RANKL-OPG system [60]. The local availability and relative preponderance of these molecules could influence the density of active osteoblasts along the bone perimeter. Mechanical gradients across an osteon and uneven availability of nutrients in the resorption cavity due for example to an off-centred capillary could also induce uneven osteoblast activity along the bone perimeter (Hypothesis H3). Delays in the initiation of bone formation on one side of the resorption cavity (Hypothesis H1) may be due to osteoblasts not yet reaching maturity there, or not being in sufficient number. We have shown in a previous study that osteoblasts need to reach a minimum density of 39/mm along the bone perimeter before they collectively form new bone [5]. We note here that our model does not include activation and inhibition mechanisms to account for these effects explicitly. These mechanisms are modelled directly through their effect on osteoblast density and activity.

The addition of asymmetry generating mechanisms in the mathematical model significantly improves model prediction (Figure 8), which suggests that osteons with lower asymmetry ratios developed by means of asymmetry generating mechanisms similar to those described by H1-H3. Our results suggest that while eclipsing lamellae may be directly related to delays in the onset of bone formation in a portion of the resorption cavity, lamellar morphology alone is unable to distinguish whether asymmetries were generated by local differences in osteoblast population, or local differences in osteoblast activity. The mathematical model shows that both are tangled in determining MAR and the morphology of successive lamellae in the osteon wall. The different results obtained under H2 or H3 in Figures 6, 7 illustrate the complex interplay between the evolution of osteoblast density and the evolution of the interface. Initially, changing either the secretory rate or osteoblast density has the same effect on the interface velocity, but these cases lead to significantly different behaviour at later times. This shows the importance of considering secretory rate and cell density independently, even if they are undistinguishable from MAR data.

Our simulations suggest that temporal cell density information could help understand what underlying mechanisms are responsible for off-centre final pores. Osteoblasts are transient cells and their density is not directly recorded in bone tissues during formation. The density of osteocytes generated in bone during formation depends on osteoblast activity rather than osteoblast density; it is determined by the ratio of secretory rate *k*_f_ and individual osteoblast embedment rate [28, 58]. Histological information on osteoblasts collected on basic multicellar units before formation has completed could provide useful information about the relative influence of osteoblast density compared to osteoblast activity in MAR and in generating osteon wall asymmetries. Osteocyte data could still provide indications of spatial heterogeneities in osteoblast numbers and osteoblast activity around a forming front.

Our study has a number of limitations:

- Our histomorphometric measurements of wall thickness asymmetry were all performed on bone biopsies from the iliac crest of women. While irregularities of bone formation in cortical osteons have been reported in a variety of skeletal sites and species, including in midshaft diaphyses of load-bearing bones [19, 21, 26, 59], it is possible that the prevalence of wall thickness asymmetry ratios is different in these other skeletal sites, particularly as the iliac crest is not load bearing. More generally, wall thickness asymmetry could depend on a number of other factors that were not considered or controlled for in our study, such as mechanical gradients, proximity of branchings with Volkmann’s canals, and irregularities of blood vessels in Haversian canals. We note here that an advantage of measuring Wall Thickness Asymmetry Ratio to quantify osteon asymmetry is that this measure is not very sensitive to section obliquity of sample preparation. A perfectly symmetric osteon with centred Haversian canal that would appear elliptic in an oblique section would still have a W.Th Asymmetry Ratio of one since the ratio of wall thickness is taken across the Haversian canal.
- Experimental data we collected in our bone samples does not contain information on osteoblasts, so osteoblast data used in the mathematical model, such as initial osteoblast density, initial secretory rates, and expected osteoblast density at completion of the osteon, are based on indirect data on MAR, secretory rates, and bone lining cell densities available in the literature [4, 43, 51].
- While the mathematical model proposed simulates bone formation within resorption cavities extracted from the histological images, it does not include biological coupling between bone resorption and bone formation [20, 60]. It assumes that bone formation can be represented accurately at a fixed cross-section of an osteon without coupling with bone resorption. The omission of bone resorption in investigations of bone formation is a common assumption [30, 32, 44], based on the spatial separation of osteoclasts and osteoblasts along the longitudinal axis of basic multicellular units. This spatial separation usually holds in the formation zone [44, 61], away from the reversal–resorption zone where osteoclasts are intermixed with pre-osteoblastic cells [5, 62]. However, in particular osteon types, such as drifting osteons, co-location of osteoclasts and osteoblasts in the same cross-section of an osteon may occur for long periods of time. The inclusion of bone resorption is necessary to model these particular osteon types.
- By considering single osteon cross-sections, our study is unable to represent three-dimensional irregularities of osteonal structures, such as spiralling osteon boundaries observed along their longitudinal axis [53, 56], and spiralling lamellae formation [59], which could be due to lateral motions of osteoblasts during bone formation [36]. Furthermore, our study did not take consideration of lamellar matrix formation motifs [63, 64]. Different lamellar organisations, such as orthogonal plywood, irregular plywood, and twisted plywood models, could reflect changes in bone formation that may induce wall thickness asymmetries. In the model, we assume that bone surface position at regular time intervals is a good representation of lamella boundaries, which is consistent with the suggestion that some lamellae correspond to a single phase of synthesis with continuous change in mineralised collagen fibril orientation [63].
- Finally, calibration of the mathematical model is based on analysing discrepancies between cortical pores generated by the mathematical model, and cortical pores measured experimentally, but not on the size and shape of the successive lamellae. Including such information at present is challenging, due to the lack of specific information about time in experimental data. Such limitation could be lifted by using fluorochrome labelling techniques and could lead to more precise, space and time dependent information on asymmetry generating mechanisms.

## 5. Conclusions

Our study reveals that asymmetry of wall thickness in secondary osteons is highly prevalent across all adult ages. This asymmetry is commonly averaged out to calculate mean osteon wall thicknesses. However, our finding that wall thickness asymmetry increases in old age indicates that this asymmetry could be an important factor in age-related bone loss and bone disorders. Future work could investigate links between wall asymmetry and other important variables of cortical osteons, including skeletal site, local mechanical variables, and quantities that have been correlated with average wall thickness, such as osteocyte lacuna density and MAR [40, 65, 66]. Such studies could provide new insights into the regulation of bone matrix deposition by osteoblasts, and how this regulation evolves with age or disease.

Most mathematical models assume symmetrical resorption cavities and symmetrical bone formation in cortical osteons [30, 61, 67, 68]. Given the prevalence of wall thickness asymmetries in cortical osteons, generalising such models to include irregular geometries is important, but also challenging as it involves modelling biological processes with strong spatial heterogeneities, and nontrivial comparisons with experimental images. The error metrics we proposed based on the position, size, and shape of the final Haversian canals provide a new systematic way to assess how well mathematical models of cortical bone formation can reproduce observed osteons. Such metrics are important to direct the development of future models into robust predictive tools.

By simulating bone formation in irregular resorption cavities extracted from experimental images, our mathematical model is able to provide new insights into bone formation processes. In particular, to generate smooth Haversian canals, our model suggests that the inevitable local geometric crowding of osteoblasts during pore filling must be finely balanced by processes helping to even out osteoblast density, such as cell–cell mechanical interactions [45–48] and geometry-dependent osteoblast elimination rate.

Our study also illustrates how mathematical modelling can be used to extract more information from experimental data and help interpret possible deregulations of bone formation. Testing various asymmetry-generating mechanisms in a mathematical model can be used as a quantitative tool to help analyse causes leading to osteons with significant asymmetries that may be associated with age or disease. Our combined experimental and mathematical study shows that we can draw links between morphological features observed in osteon cross-sections, and specific biological mechanisms leading to asymmetric wall thicknesses during cortical pore filling. However, our results also suggest that it may not always be possible to disentangle the nature of some asymmetry-generating processes without more detailed information on osteoblast densities and their activity. Gaining deeper insights into these mechanisms would help the calibration, validation and predictive capabilities of future mathematical models. These insights are also particularly important given our finding that wall asymmetry ratio evolves with age, as it could lead to more detailed diagnoses of specific disruptions of bone formation from morphological observation of osteons in bone samples.

## Acknowledgments

This research is supported by the Australian Research Council (DP190102545, DP230100025) and the Centre for Biomedical Technologies, Queensland University of Technology (QUT). We thank Dr Mohd Almie Alias for sharing computer code developed for Ref. [34].

## A. Mathematical model

All the equations of the mathematical model used to perform the numerical simulations are summarised in this appendix. For their derivation and justification, the reader is referred to Refs [34, 36].

### A.1 Evolution of osteoblast density and of osteonal bone perimeter

The evolution of the osteonal bone perimeter during formation is entirely specified by the normal velocity

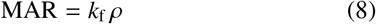

at each point of the bone perimeter. This velocity depends on both time and location, and on the density of osteoblasts. The evolution of osteoblast density is governed by the following partial differential equation [33, 36]:

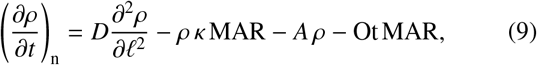

where *ℓ* is an arc length coordinate around the current bone perimeter, *κ* is the signed curvature of the bone perimeter (*κ <* 0 where concave, *κ >* 0 where convex), and ^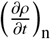^ represents the time rate of change of osteoblast density in the normal direction to the interface [36].

In circular osteons, experimental observations suggest that *A* and *k*_f_ depend on pore radius, *R* [34, 43]. When osteons are not circular, the dependence on *R* represents an influence of pore curvature *κ* for the elimination rate of osteoblasts, and an influence of pore area *ϕ* for the secretory rate [34], such that

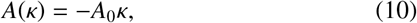

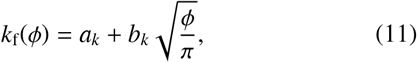

where *a*_*k*_ = 3.2741 ×10^−6^ mm^2^*/*day, and *b*_*k*_ = 8.5728 ×10^−5^ mm^2^*/*day [34].

### A.2 Asymmetry generating mechanisms

#### Delayed formation

A delayed onset of formation in a portion of the bone perimeter is modelled by setting osteoblast elimination rate and osteoblast secretory rate to zero in this portion of the perimeter for a given amount of time *T* . This is achieved by multiplying the expressions in Equations (10),(11) by a switch function *c*_delay_(*t, θ*), as follows:

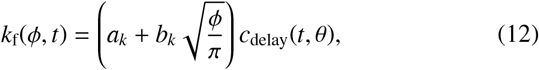

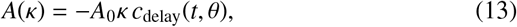

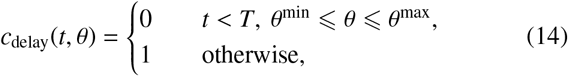

where *θ* is a polar angle in the plane of the osteon cross-section. The origin of the coordinate system is placed at the centroid of the osteon pore in the experimental images.

#### Uneven osteoblastogenesis

A nonuniform initial distribution of osteoblasts is modelled by modulating the initial osteoblast density *ρ*_0_ = 161 cells mm^−1^ with a varying function of the polar angle:

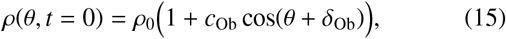

where 0 *< c*_Ob_ *<* 1 is the amplitude of the perturbation, and *δ*_Ob_ is an angular shift. Values assumed for these parameters are osteon-specific and mentioned in the figure captions.

#### Uneven secretory rate

Uneven osteoblast activity around the bone perimeter is modelled by modulating secretory rate with a varying function of the polar angle:

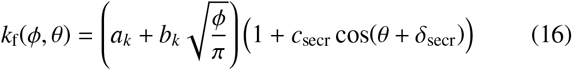

where 0 *< c*_secr_ *<* 1 is the amplitude of the perturbation, and *δ*_secr_ is an angular shift.

### A.3 Numerical discretisation

The mathematical model describing the co-evolution of osteoblast density and interface position in Eqs (8),(9) is solved numerically using a Kurganov–Tadmor finite volume method [69] based on computer code developed by M. A. Alias [33, 34]. The numerical scheme solves for the position of the interface and osteoblast density along the interface in polar coordinates in conservative form, see Ref. [33] for further details. This numerical scheme is chosen for its conservation properties and high resolution, which are particularly important due to the small-scale irregularities of the initial interfaces extracted from the experimental images. The polar angle is discretised using 120 points in all simulations (corresponding to an angular spatial step Δ*θ* = 3^*°*^). The system of ordinary differential equations obtained from the semi-discrete Kurganov–Tadmor scheme after spatial discretisation is solved using a third order Runge–Kutta scheme [70] with a time step Δ*t* adapted to the value of diffusivity chosen:

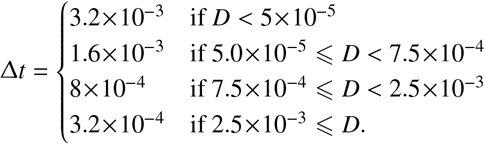

Initial conditions assume a constant density of osteoblasts *ρ*_0_ = 161 cells mm^−1^ (Table 1) except in simulations using the asymmetry-generating Hypothesis H2 (uneven osteoblastogenesis) where the initial density is given by Eq. (15). The initial interface is the osteon resorption cavity (cement line) extracted from the experimental images. Periodic boundary conditions are used on the polar angle.

## B. Calculation of error metrics

The error metrics in Eqs (2)–(6) are calculated using the following formulae.

The average osteoblast density 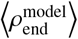 in Eq. (2) is defined as the line integral [71]

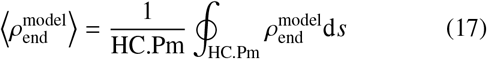

where HC.Pm is the perimeter of the Haversian canal. This integral is calculated based on the polygonal discretisation of the Haversian canal boundary.

Haversian canal pore areas in Eq. (3) are defined as double integrals over the Haversian canal region *R*

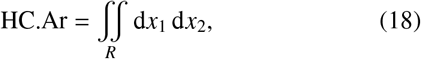

and calculated numerically using the so-called surveyor’s formula or shoelace formula from a polygonal discretisation of the Haversian canal boundary [71].

Haversian canal centroids in Eq. (4) are the geometric mean of the Haversian canal region, given by the first moment of the double integral over *R*. If ***r*** = (*x*_1_, *x*_2_), then

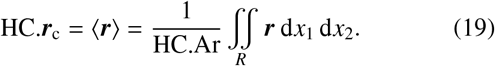

The orientation *θ* and aspect ratio *a/b* of the ellipse that best fit a Haversian canal region in Eqs (5),(6) are obtained by calculating second moments of the Haversian canal region:

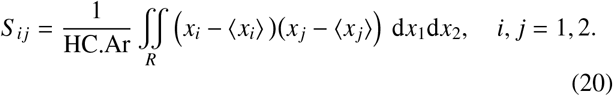

The covariance matrix *S* = {*S* _*i j*_} _*i, j*=1,2_ that these second moments define is real and symmetric, and defines the ellipse that best fits the Haversian canal region *R* via the equation

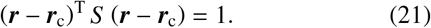

This ellipse is centred at the centroid ***r***_c_. The values and directions of the ellipse’s semi-axes are given by the eigenvalues and corresponding eigenvectors of *S*, respectively. The orientation *θ* of the ellipse with respect to the *x* axis is given by

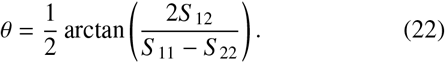

The ratio of eigenvalues of *S* is the ratio *a/b* of the ellipses’ largest over smallest semi-axes, given by

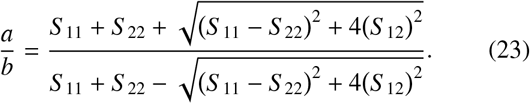

